# A CRISPRi Library Screen in Group B *Streptococcus* Identifies Surface Immunogenic Protein (Sip) as a Mediator of Multiple Host Interactions

**DOI:** 10.1101/2024.12.06.627252

**Authors:** K Firestone, KP Gopalakrishna, LM Rogers, A Peters, JA Gaddy, C Nichols, MH Hall, HN Varela, SM Carlin, GH Hillebrand, EJ Giacobe, DM Aronoff, TA Hooven

**Author notes:** Authors contributed equally (K Firestone, KP Gopalakrishna).

## Abstract

Group B *Streptococcus* (GBS; *Streptococcus agalactiae*) is an important pathobiont capable of colonizing various host environments, contributing to severe perinatal infections. Surface proteins play critical roles in GBS-host interactions, yet comprehensive studies of these proteins’ functions have been limited by genetic manipulation challenges. This study leveraged a CRISPR interference (CRISPRi) library to target genes encoding surface-trafficked proteins in GBS, identifying their roles in modulating macrophage cytokine responses. Bioinformatic analysis of 654 GBS genomes revealed 66 conserved surface protein genes. Using a GBS strain expressing chromosomally integrated dCas9, we generated and validated CRISPRi strains targeting these genes. THP-1 macrophage-like cells were exposed to ethanol-killed GBS variants, and pro-inflammatory cytokines TNF-α and IL-1β were measured. Notably, knockdown of the *sip* gene, encoding the Surface Immunogenic Protein (Sip), significantly increased IL-1β secretion, implicating Sip in caspase-1-dependent regulation. Further, Δ*sip* mutants demonstrated impaired biofilm formation, reduced adherence to human fetal membranes, and diminished uterine persistence in a mouse colonization model. These findings suggest Sip modulates GBS-host interactions critical for pathogenesis, underscoring its potential as a therapeutic target or vaccine component.

## Background and Introduction

*Streptococcus agalactiae* (group B *Streptococcus*; GBS) is an encapsulated, gram-positive pathobiont that asymptomatically colonizes the intestine and reproductive tracts of approximately one-third of healthy adults, but also causes opportunistic infections, particularly during pregnancy, the neonatal period, and infancy^1,2^. GBS exhibits niche versatility in the human host, persisting in the intestinal lumen^3,4^, the vagina^5–7^, within the pregnant uterus (including placental tissue, fetal membranes, amniotic fluid, and the fetus)^8,9^, the newborn bloodstream^10–12^, and within cerebrospinal fluid^13,14^. This versatility, and particularly the ability to evade innate and adaptive immune clearance in anatomically and immunologically protected gestational compartments, contributes to GBS pathogenicity during the perinatal period.

GBS persistence during interactions within diverse host environments is mediated by bacterial surface features. The GBS sialylated polysaccharide capsule, of which there are ten known subtypes defined by their patterns of molecular cross-linkage, has been shown to play key roles in immune evasion and subversion. The GBS capsule promotes biofilm formation and epithelial colonization^15^, and influences cytokine responses by leukocytes after surface contact^16^, among other roles.

Within and extending beyond the GBS capsule are surface-anchored and secreted proteins. Like the polysaccharide capsule, some externalized GBS proteins are known to promote fitness in otherwise unhospitable host environments. GBS pilus proteins enable host surface attachment^17,18^ and biofilm formation^19^. HvgA is an adhesin whose roles in promoting neonatal intestinal adhesion, transmural invasion, and attachment to and passage across the blood-brain barrier are well-described^13,20^. The serine repeat proteins (Srr1 and Srr2) are important adhesins whose roles in perinatal GBS pathogenesis are also well-characterized^21–23^. C5a peptidase contributes to immune evasion by cleaving complement whose surface deposition aids phagocytotic clearance^24^, and plays a moonlighting role as an adhesin^25–27^. Other surface-trafficked proteins include sensor and signal transduction proteins that bind to and relay detection of diverse environmental solutes^28,29^. Another large class of surface-trafficked GBS proteins are those involved in chemical flux into and out of the cell.

GBS employs multiple genetically encoded trafficking motifs to direct proteins to the cell surface, move them across the cell membrane, and either anchor them in place or secrete them into the external environment. Signal peptide sequences, encoded at the N- termini of surface-trafficked proteins, interact with components of the bacterial Sec system, which recognize signal peptide-containing proteins, chaperone them to and across the bacterial surface, then cleave and degrade the signal peptide trafficking flag^30^. Signal peptide sequences often co-occur with surface anchoring motifs, the most common of which in GBS is LPXTG^31^. These motifs interact with sortase enzymes whose role is to orient and attach a subset of surface-trafficked proteins to the GBS cell wall exterior^31^.

While numerous GBS surface-trafficked proteins have been studied and described, obstacles have limited large-scale and systematic examination of their function. One important challenge has been the limited ability to perform high-throughput, targeted genetic manipulation. Traditional approaches to GBS mutagenesis rely on double- crossover allelic exchange techniques that are inefficient and prone to creating unintended rearrangements^32,33^.

CRISPR interference (CRISPRi) is an alternative to generation of chromosomal mutants for studying curated gene sets. Rather than creating and validating individual gene knockouts, which is throughput-limiting in GBS, CRISPRi leverages a catalytically inactive Cas protein (dead Cas; dCas) to sterically block transcription at a specific genomic locus^34^. The major advantage of CRISPRi over traditional mutagenesis approaches is that the targeting portion of the single guide RNA (sgRNA) sequence can easily be changed by encoding it on a modular plasmid. This allows targeted alteration of gene expression following a few short cloning and transformation steps.

We recently introduced a system for creating CRISPRi gene knockdown strains in GBS^35^. Our strategy uses a GBS mutant background in which two point mutations convert wild type (WT) GBS Cas9 to dCas9, expressed from the chromosome at its native locus. Into this dCas9-expressing background we introduce a modular sgRNA encoded on a shuttle vector, p3015b. A series of straightforward recombinant DNA reactions using custom ordered oligonucleotides allows rapid reprogramming of dCas9 to target genetic loci on the chromosome. In our initial publication about the GBS CRISPRi system, we confirmed that changing WT Cas9 to dCas9 does not have significant off-target effects on gene expression^35^. Therefore, phenotypic effects of dCas9-mediated gene knockdown can be presumed to arise from the targeted gene.

In this study, we turn from GBS CRISPRi proof-of-concept to using the technology to create and study a curated library of targeted knockdown strains. Because of the importance of externalized proteins in host and environmental interactions, we aimed to generate and examine a knockdown library comprising a large set of surface-trafficked proteins. We identified targets by the presence of signal peptide sequences encoded at their N-termini and screened a set of over 600 GBS genomes to establish which signal peptide encoding genes were conserved across this large collection of isolates. Because the effects of most GBS surface proteins on innate immune cell responses are unknown, we opted to screen the library we created for altered cytokine triggering effects on cultured macrophage-like THP-1 cells^36^.

We found that altering GBS surface protein expression by CRISPRi had considerable effects on THP-1 macrophage release of pro-inflammatory cytokines TNFα and IL-1β. While some surface protein knockdown strains led to decreased cytokine release, a substantial portion of the knockdown strains in our library led to increased cytokine expression. A knockdown strain targeting the highly conserved GBS surface immunogenic protein (Sip) gene led to the greatest increase in IL-1β release and significantly increased TNF-α release from THP-1 cells. Recombinant Sip has been previously described as a potential GBS vaccine component and tested in animal models of prematurity and GBS infection^37–41^.

However, Sip’s role in GBS biology and its interactions with host cells and surfaces has not previously been described.

We elected first to focus on the IL-1β response, as mature IL-1β release is one of the culminating processes of NLRP3 inflammasome activation, which is recognized as a key factor in triggering preterm labor and stillbirth in pregnancies affected by bacterial chorioamnionitis or sterile inflammation^42–45^. NLRP3 inflammasome activation leads to caspase-1 cleavage of pro-IL-1β, generating mature IL-1β that is then secreted through gasdermin-D pores, whose presence is also caspase-1 dependent^46^. Using in-frame *Δsip* knockout mutants in two GBS strains and testing in several *in vitro*, *ex vivo*, and *in vivo* models of GBS colonization and disease, we examined Sip’s significant influence on IL-1β transcription and caspase-1-mediated post-translational processing. Unexpectedly, we also found that Sip plays a significant role in GBS biofilm formation and association with host gestational tissues. Together, our results suggest that Sip influences multiple bacterial-host interactions implicated in pathogenesis of perinatal complications.

## Results

### Bioinformatic identification of conserved GBS surface-trafficked proteins

To develop a set of target genes with N-terminus signal peptide sequences, we used publicly available GBS genomes and bioinformatic tools provided by the United States Department of Energy Joint Genome Institute’s (JGI) Integrated Microbial Genomes and Microbiomes System (IMG/M; https://img.jgi.doe.gov/m/)^47–49^. IMG/M’s microbial genome annotation pipeline includes a built-in signal peptide designation for genes and allows gene sequence conservation analysis across a large user-defined set of genomes.

First, we used the signal peptide search criterion to extract the complete set of signal peptide genes from the genome of CNCTC 10/84, the strain background in which we planned to generate CRISPRi gene expression knockdowns. Next, using *Streptococcus agalactiae* as the species designator, we generated a set of 654 genomes posted to the server at that time. Then we used the Gene Profile function on IGM/M to quantify unidirectional sequence similarity for the complete set of signal peptide genes among our set of 654 GBS genomes using a maximum E-value of 0.1. This generated a set of 75 genes that were present in CNCTC 10/84 and in at least a subset of our GBS genome screening set. The mean sequence conservation in the set was 80% with standard deviation 21%. The wide standard deviation was driven by nine genes with less than 50% homology across the genome collection. Excluding these low-homology genes brought conservation among the set to 88% with standard deviation 5.2%. The 66 genes in the gene set excluding the low-homology matches were considered the set of conserved surface-trafficked protein genes (**Fig. 1A**, **Sup. Data 1**). We then used SignalP server to analyze the conserved surface- trafficked gene sequences, generating positional function predictions for the N-termini of all genes in the set^50^. This analysis confirmed predicted presence of signal peptide sequences at the start of our target gene set (**Fig. 1B**).

**Figure 1:**
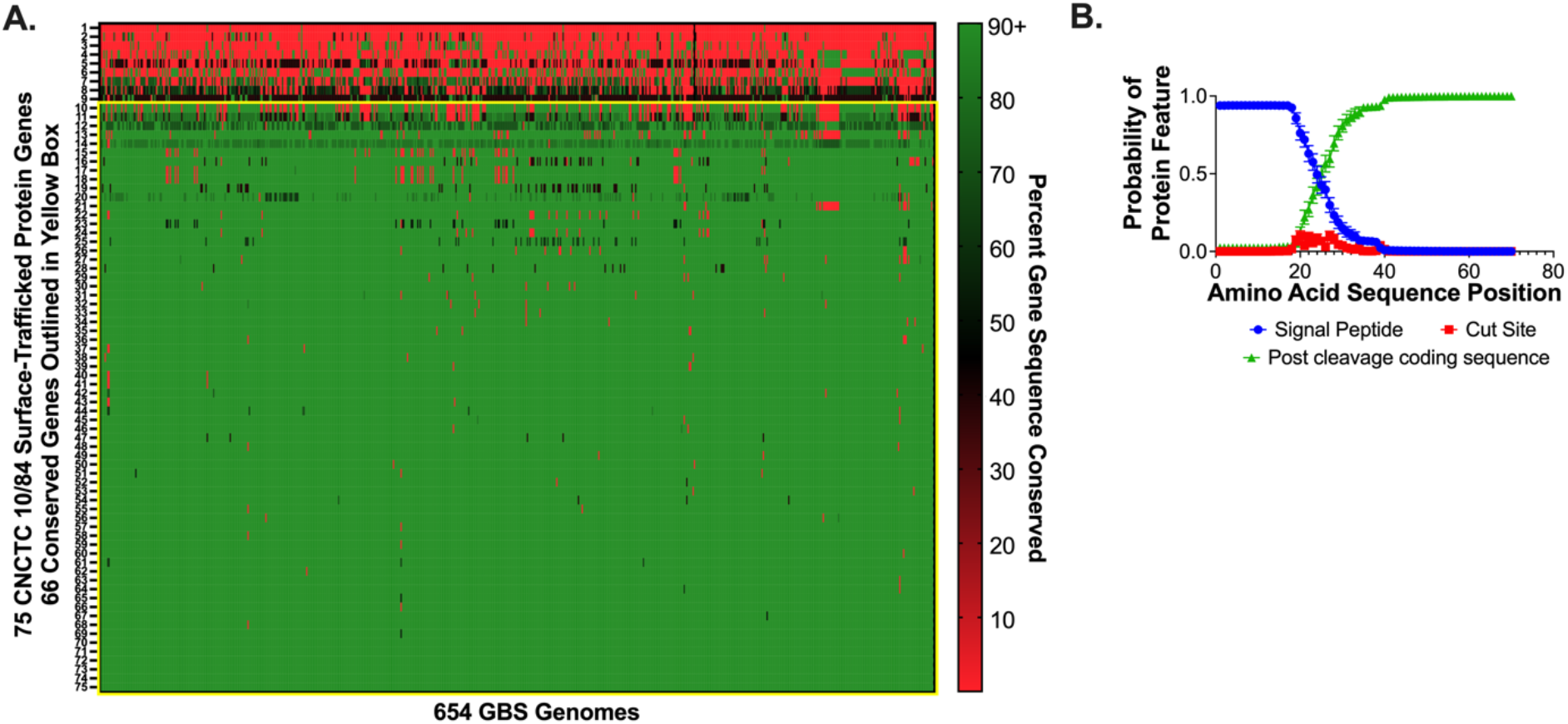
Bioinformatic identification and verification of conserved signal peptide- encoding GBS genes. A heat map shows percent gene sequence conservation among 75 CNCTC 10/84 genes with signal peptide sequences, when compared across 654 GBS genomes posted to the Integrated Microbial Genomes and Microbiomes System. The genes outlined in the yellow box are the set 66 of high-homology genes used as the conserved surface-trafficked gene set for CRIPSRi library generation (**A**). SignalP verification of N-termini signal peptide-encoding sequence motifs in the CRISPRi gene set (**B**). The mean probability of signal peptide sequence features (signal peptide, signal peptide cut site, post-cleavage coding sequence) is graphed by amino acid position for the conserved surface protein set. Error bars show standard error of the mean.

### Generation and validation of a CRISPRi library targeting the set of signal peptide genes

Using our previously described cloning process for CRISPRi^35^, we sought to generate CRISPRi knockdowns targeting the conserved set of CNCTC 10/84 surface-trafficked proteins. The GBS protospacer adjacent motif (PAM), required for dCas9 binding to chromosomal DNA, is NGG, which is identical to spyCas9 from *Streptococcus pyogenes*^51–53^. We therefore used the publicly available CRISPick server with *S. pyogenes* settings to analyze gene target sequences and rank potential targeting protospacers^54,55^. We filtered for sgRNA protospacer sequences complementary to the antisense strand of each gene, as this is necessary for optimal steric hinderance of RNA polymerase and gene knockdown^34,35^. Whenever possible, we selected CRISPRi targets in the leading one third of a gene’s coding sequence, since dCas9 binding to the coding sequence terminus leads to decreased knockdown efficacy. Our goal was to make a CRISPRi library with two sgRNA targets per gene. This redundancy was because we and others have observed that sgRNA efficiency is variable in CRISPRi systems, even when design criteria, such as outlined above, are employed.

Following protospacer design, we obtained corresponding custom oligonucleotides, which we annealed to generate double-stranded inserts for cloning into *BsaI*-digested expression plasmid p3015b, followed by transformation into chemically competent DH5α *Escherichia coli*. Transformant colonies were tested for correct cloning of the intended protospacer using colony PCR in which one primer was the forward oriented protospacer oligonucleotide, paired with a reverse primer complementary to p3015b downstream of the protospacer insertion site. Using this approach, only colonies with the correct protospacer insert would generate a PCR product, which was visualized by agarose gel electrophoresis.

Correctly recombinant p3015b clones with the intended protospacer sequences were used for plasmid miniprep. Miniprepped plasmid was then used to transform electrocompetent GBS strain CNCTC 10/84 bearing *dCas9* on its chromosome (10/84:*dcas9*). Colonies from this transformation were grown overnight and stored as frozen stocks. Some of our intended knockdown strains could not be completed after several attempts, either because the sgRNA cloning step failed in *E. coli*, or the apparently successfully generated plasmid could not be transformed into electrocompetent 10/84:*dcas9*, or colonies of transformed 10/84:*dcas9* had little to no growth in liquid culture. Five genes among the 66 conserved surface protein genes were not targetable.

Most of the remaining 61 were targeted with two protospacers; 16 were targeted with a single protospacer.

The 106 successfully transformed 10/84:*dcas9* variants, each bearing a unique protospacer targeting a member of the surface-trafficked gene set, were then used for reverse transcriptase quantitative polymerase chain reaction (RT-qPCR) to test expression of the targeted gene in each knockdown strain. For this testing, expression of the recombinase gene *recA* was used as an RNA normalization control. Individual knockdown strains were grown overnight and assayed in triplicate biological replicates. Sham-targeted 10/84:*dcas9* was used as the control comparator for each gene.

Most of the knockdown strains in our collection were downregulated. As has been observed in other CRISPRi systems^55–57^, a wide range in degree of knockdown was observed, spanning from almost undetectable levels (2^-12^) to 13 strains that had equivalent or even increased expression relative to the nontargeted control. Mean expression knockdown across the entire collection was 0.365 of control (**Fig. 2**, **Sup. Data 1**).

**Figure 2:**
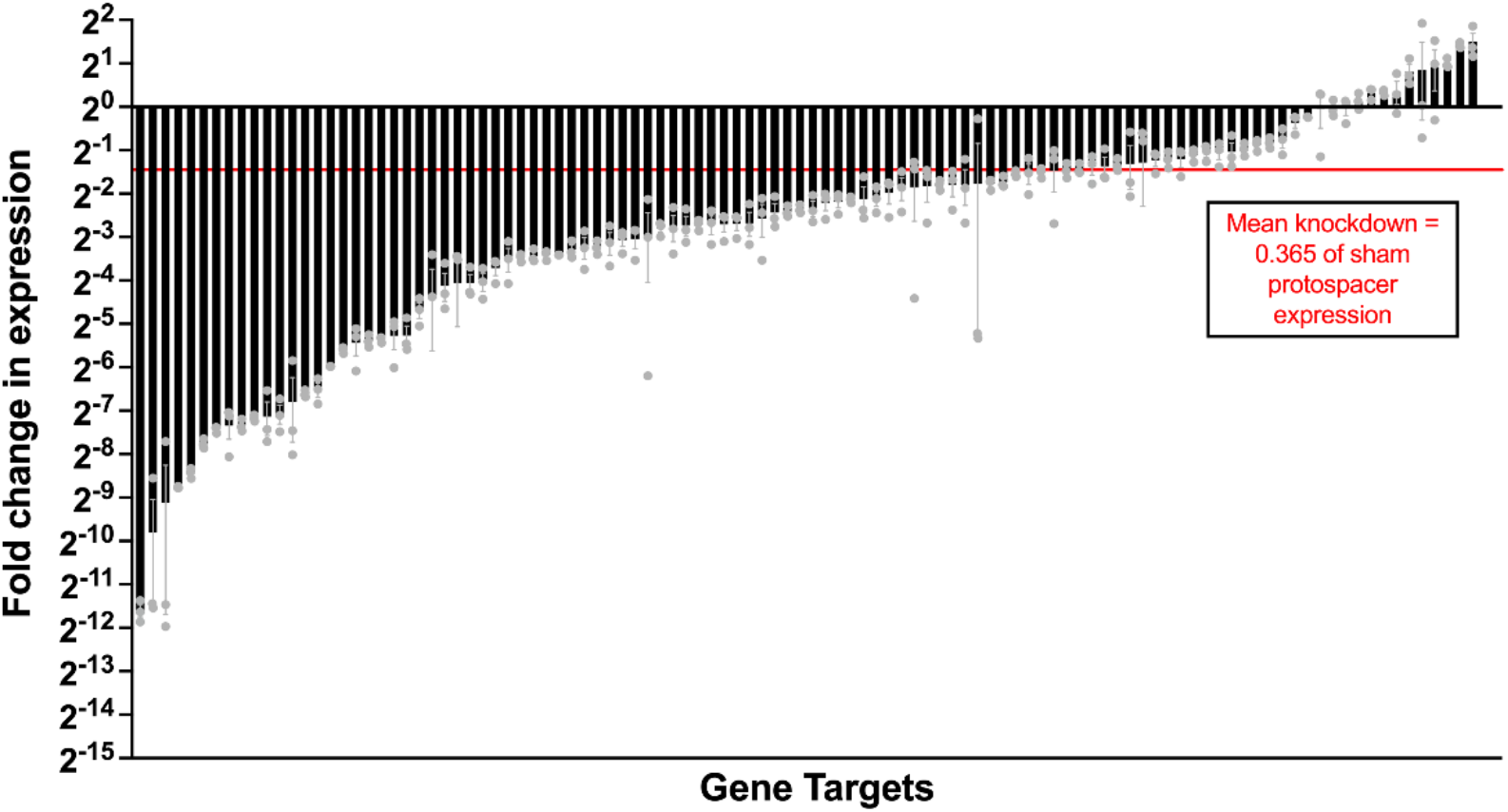
RT-qPCR validation of conserved signal trafficked protein knockdown library. Triplicate independent biological replicates of 106 CRISPRi strains from the conserved surface protein knockdown library were grown and used for RNA extraction. RT-qPCR was performed to compare target gene expression to a sham-targeted isogenic control strain of CNCTC 10/84:*dcas9* (RNA concentration was normalized to the housekeeping gene *recA*). Error bars in the figure show mean fold-change values with error bars delineating standard error of the mean.

### Screen for cytokine responses by THP-1 macrophage-like cultured cells

To examine phagocyte pro-inflammatory responses to GBS with suppressed surface protein expression, we screened 103 knockdown variants, in triplicate biological replicates, for THP-1 cell induction of the high-level cytokines TNF-α and IL-1β after they had been differentiated to a macrophage phenotype with phorbol 12-myristate 13-acetate (PMA; **Fig. 3A**). For this testing, we excluded 20 GBS variants that had not met the knockdown criterion of <0.5 control expression of their target gene. Because of the possibility that differences in growth rate, intracellular survival, or macrophage killing capacity among our variants might confound our measures, and because the genes we had targeted by CRISPRi were selected based on the exteriority of their protein products, we used standardized, ethanol-killed preparations of the variants and controls in our GBS collection, rather than live bacteria.

**Figure 3:**
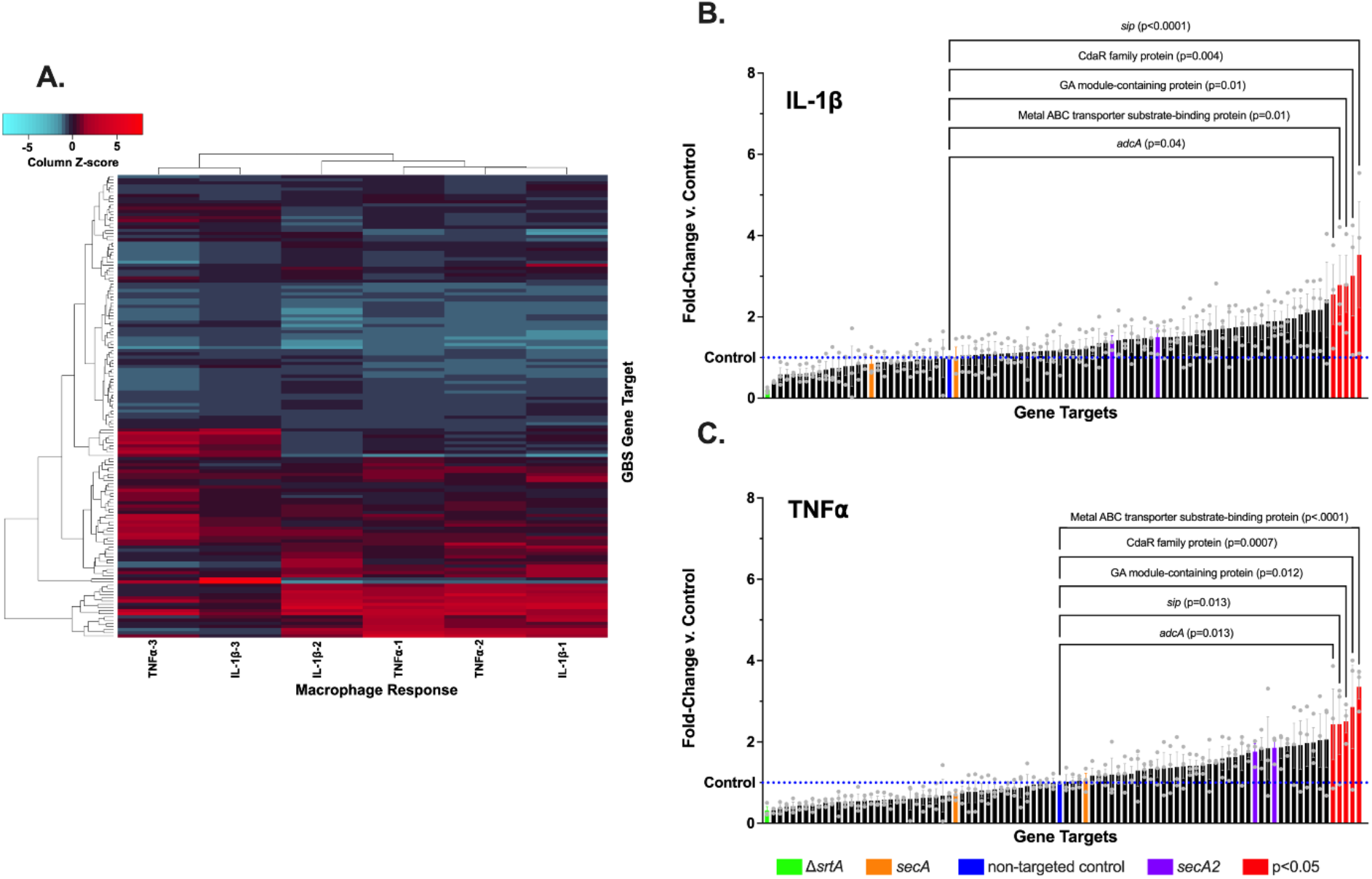
THP-1 macrophage cytokine responses to GBS from the conserved surface trafficked protein CRISPRi library. THP-1 cells were exposed to ethanol-killed knockdown strains from the CRISPRi library and assayed using ELISA for IL-1β and TNFα. Each exposure/assay pair was performed in independent biological triplicates. The heatmap (**A**) shows column-normalized, hierarchically clustered data from the CRISPRi library strains. IL-1β (**B**) and TNFα (**C**) data are shown with inclusion of four control strains included in the experiment (Δ*srtA*, *secA*, sham, *secA2*). Statistically significant (*p<0.05*) comparisons to sham with one-way ANOVA and FDR correction (Q=0.05) are labeled red. Each data point represents an independent biological replicate (n=3); error bars show standard error of the mean.

We included five additional strains in these experiments with genomic targeting of chaperones involved in surface protein localization. The first was a CNCTC 10/84 Δ*srtA* knockout with a chromosomal deletion of the sortase A gene. As mentioned above, the SrtA enzyme is necessary for proper attachment of LPXTG-containing proteins to the outer surface of the GBS cell wall^31^. We also included CRISPRi knockdowns, two each, of the *secA* and *secA2* genes. SecA is the major effector of signal peptide-bearing protein translocation across the cell membrane. SecA2, by contrast, is a more specialized surface protein chaperone likely involved in shuttling a subset of GBS surface-trafficked proteins, including the glycosylated serine repeat adhesin Srr1^58^.

Statistical testing of mean TNF-α and IL-1β measurements across all three biological replicates was performed between all bioengineered variants and CNCTC 10/84:*dcas9* transformed with the sham, nontargeting p3015b plasmid, using one-way ANOVA with a Q=0.05 false discovery rate (FDR) correction for multiple comparisons. This approach yielded five knockdowns with above-threshold changes in the IL-1β assay (**Fig. 3B**, **Sup. Data 1**) and five knockdowns with above-threshold changes in the TNF-α assay (**Fig. 3C**, **Sup. Data 1**). Interestingly, the discoveries were all among knockdowns that resulted in increased cytokine expression by the THP-1 macrophages following coincubation. Although the mean cytokine expression was decreased after coincubation with a subset of our knockout collection, none of these decreased cytokine conditions met prespecified criteria for statistical significance.

The knockdown variants with statistical significance in both assay sets are listed in **Table 1**. There was full concordance for strains appearing in the statistically significant subset for both cytokine assays.

**Table 1:** GBS genes with significant eRects on THP-1 cell cytokine secretion Signal peptideencoding gene annotation CNCTC 10/84 Refseq locus Protospacer target nucleotide Mean IL-1β fold-change vs. control strain (pvalue) Mean TNF.

| **Table 1: GBS genes with significant effects on THP-1 cell cytokine secretion** |  |  |  |  |
| --- | --- | --- | --- | --- |
| **Signal peptide-encoding gene annotation** | **CNCTC 10/84 Refseq locus** | **Protospacer target nucleotide** | **Mean IL-1β fold-change vs. control strain (p-value)** | **Mean TNFα fold-change vs. control strain (p-value)** |
| *sip* | W903_RS00330 | 271 | 3.53 (<0.0001) | 2.44 (0.013) |
| CdaR family protein | W903_RS04760 | 325 | 3.01 (0.004) | 2.86 (0.0007) |
| GA module-containing protein | W903_RS04495 | 294 | 2.81 (0.01) | 2.51 (0.012) |
| Metal ABC transporter substrate-binding protein | W903_RS07525 | 199 | 2.78 (0.01) | 3.36 (<0.0001) |
| *adcA* | W903_RS03470 | 199 | 2.55 (0.04) | 2.43 (0.013) |

### Targeted Sip deletion strains lead to increased phagocyte IL-1β secretion in a caspase-1 dependent manner

Among the conserved, targeted, surface-trafficked proteins whose CRISPRi repression had a significant effect on THP-1 cytokine release, our knockdown of the surface immunogenic protein (Sip) gene had the greatest effect on IL-1β. This gene was interesting to us for several reasons. First, mature IL-1β has important roles in triggering preterm labor and stillbirth in pregnancies affected by bacterial chorioamnionitis or sterile inflammation, as described in the introduction. Second, Sip has been examined in several studies as a candidate recombinant protein vaccine. Preparations of Sip have shown protective effects in animal models of vaginal colonization and neonatal sepsis^38,59,60^. However, no studies of Sip have focused on the protein’s role in its natural setting as a GBS surface factor.

Therefore, to examine the bacterial cell biology of Sip, we generated in-frame deletion knockouts of the *sip* gene in two GBS background strains: CNCTC 10/84, the same serotype V strain we had used for our CRISPRi screen, and A909, a serotype Ia strain first collected from a septic neonate.

To make Δ*sip* deletion mutants in these two background strains, we used a temperature- and sucrose-based counterselection mutagenesis plasmid, pMBsacB^32^. The final step in allelic replacement mutagenesis with pMBsacB is a crossover event, in which the plasmid excises from the chromosome through recombination between homologous DNA sequences upstream and downstream of the target gene. This recombination event can either lead to a deletion mutant or reversion to wild type (*rev*). After performing PCR to confirm mutant and revertant genotypes in different GBS clones that resulted from genetic manipulation, we whole genome sequenced CNCTC 10/84 and A909 Δ*sip* and *rev* strains, determining that they did not harbor potentially confounding off-target mutations. In experiments examining phenotypes of the Δ*sip* mutants, we used the corresponding *rev* strains—which had been through all the same outgrowth phases, temperature shifts, and sucrose exposures as the knockouts—as a set of controls.

We assessed ethanol-killed CNCTC 10/84 and A909 Δ*sip* and *rev* strains in coincubation experiments with THP-1 cells, testing by ELISA for secretion of IL-1β (**Fig. 4A**) and TNF-α (**Fig. 4C**) as we did in our initial screen. Consistent with the screen results, both 10/84:Δ*sip* and A909:Δ*sip* demonstrated increased induction of THP-1 macrophage IL-1β release into the supernatant, relative to revertant strains (**Fig. 4A**). There was also an increase in TNF-α release during A909:Δ*sip* coincubation compared to A909:*rev* (**Fig. 4C**). A slight increase in TNF-α following THP-1 coincubation with 10/84:Δ*sip* was measured, but was statistically nonsignificant (**Fig. 4C**).

**Figure 4:**
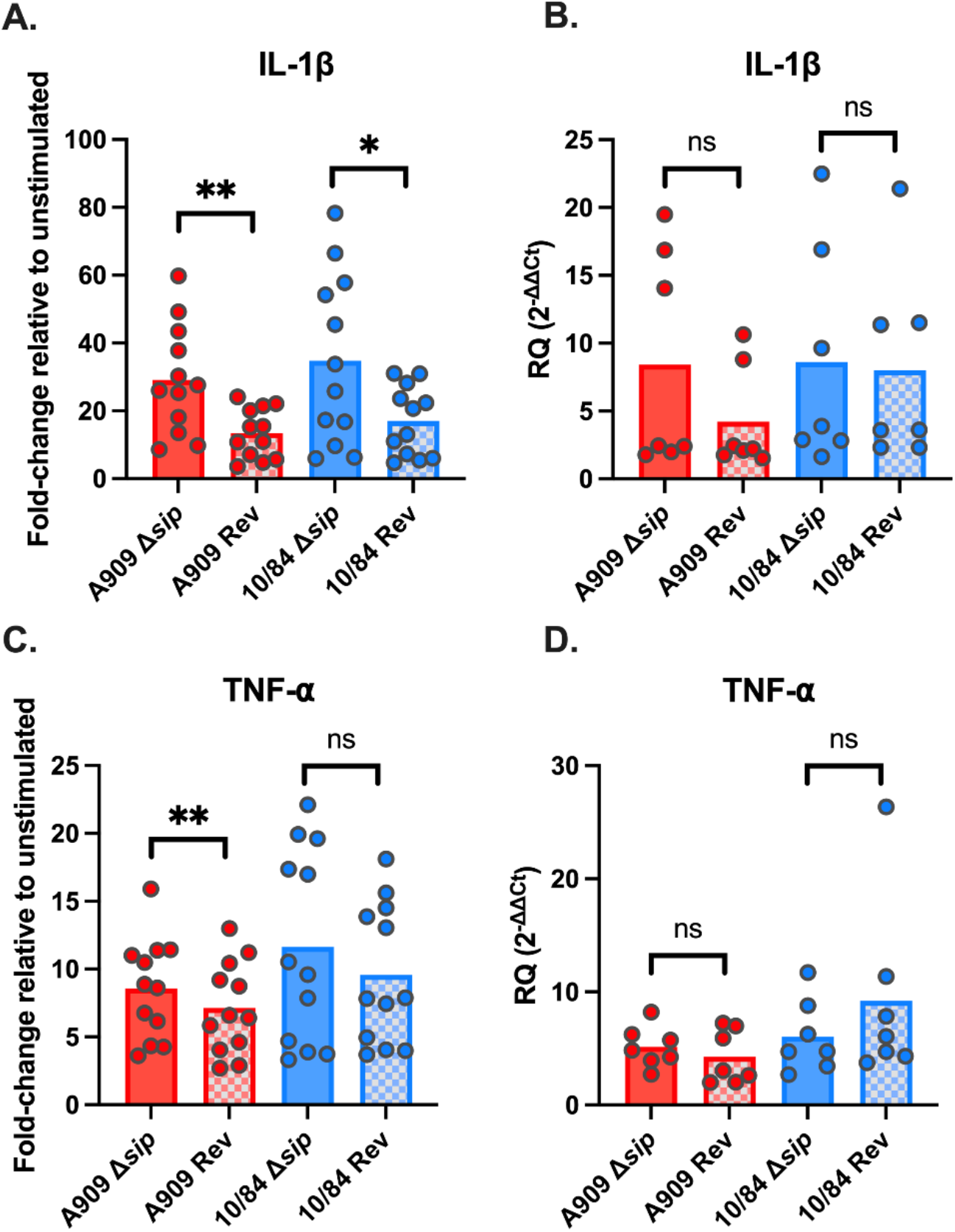
THP-1 macrophage cytokine responses to targeted Sip deletion mutant GBS strains. THP-1 cells were exposed to ethanol-killed Δ*sip* or revertant (*Rev*) strains using a 50:1 MOI. IL-1β (**A-B**) and TNFα **(C-D)** cytokine responses were measured by ELISA (**A, C**, n=12) or RT-qPCR (**B, D**, n=7; RQ=relative quantity, Ct=Cycle threshold)). Each data point represents an independent biological replicate of THP-1 cells, which were exposed to each of the four strains in separate wells. Statistical testing by paired T-test with Bonferroni’s correction for multiple comparisons (\**p<0.05*, \*\**p<0.01*).

In addition to ELISA-based detection of IL-1β and TNF-α in the THP-1 supernatant, we used RT-qPCR to evaluate transcriptional level changes in THP-1 cells following GBS coincubation. This testing showed no differences between Δ*sip* and *rev* strains in induction of IL-1β or TNF-α gene transcription (**Fig. 4B, D**). Increased secretion of mature IL-1β without increased transcription is characteristic of the NLRP3 inflammasome response to toll-like receptor stimulation. During NLRP3 inflammasome-driven pyroptosis, pro-IL-1β present in the cytosol is cleaved by activated caspase-1 following assembly of ASC-NLRP3 complexes. Mature IL-1β is then secreted through gasdermin-D pores, whose presence is also secondary to caspase-1 activation^46^. To follow up on these findings, we introduced Z-YVAD-FMK, an irreversible caspase-1 inhibitor^61^, which led to suppression of the IL-1β stimulatory effect observed during coincubation with the Δ*sip* and *rev* strains (**Fig. 5**).

**Figure 5:**
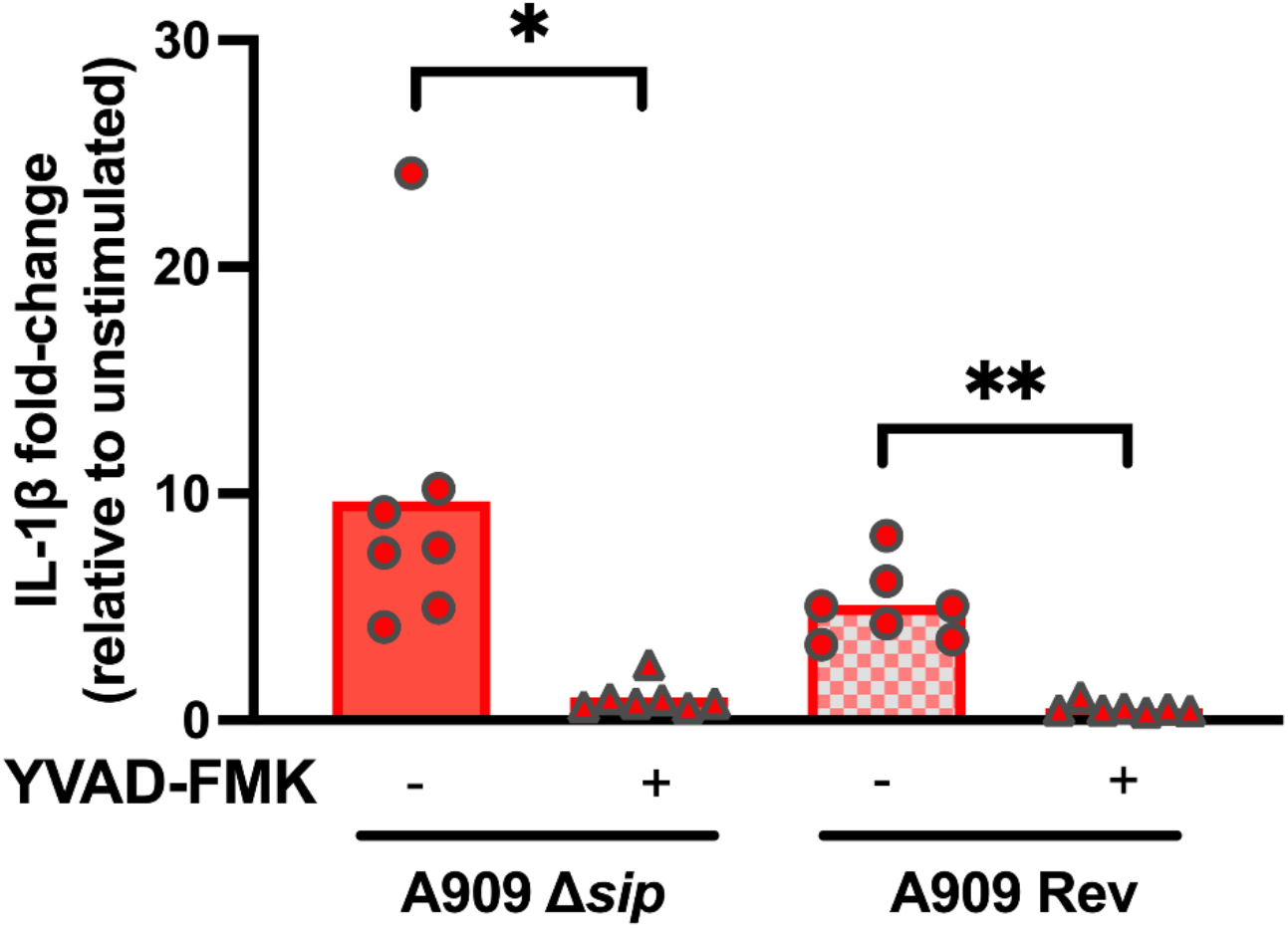
IL-1β release from GBS-exposed THP-1 macrophages is caspase-1 dependent. THP-1 cells were exposed to ethanol-killed A909 Δ*sip* or *Rev* strains, either in the presence of YVAD-FMK caspase-1 inhibitor or vehicle control. Each datapoint represents an independent biological replicate of THP-1 cells that were exposed to the experimental strains. Statistical testing by paired T-test with Bonferroni’s correction for multiple comparisons (\**p<0.05*, \*\**p<0.01*).

Together, these findings suggest that GBS Sip has an inhibitory effect on caspase-1- mediated secretion of IL-1β such that its deletion from the GBS genome leads to an increased pyroptotic response from THP-1 macrophage-like cells.

### GBS Δsip mutants are deficient at forming biofilms in vitro

GBS can form biofilms *in vivo* and *in vitro*^15,62,63^. Biofilms are thought to be a mechanistic factor in colonization of the vaginal epithelium, which in turn increases the risk of vertical transmission. Bacterial surface-trafficked proteins can affect the propensity to form biofilms, so we performed *in vitro* modeling of biofilm formation in our Δ*sip* and *rev* strains.

In quantitative colorimetric assays of biofilms grown in microtiter plates, we measured significantly less biofilm (normalized to planktonic biomass in the overnight culture) in CNCTC 10/84:Δ*sip* and A909:Δ*sip* relative to *rev* and WT controls. 10/84:Δ*sip* exhibited a 38% attenuation in its ability to form biofilms compared to the parental strain and a 41% attenuation when compared to the 10/84:*rev* control (*P<0.05, one-way ANOVA with Tukey’s *post hoc* correction). A909:Δ*sip* exhibited a 59% attenuation in its ability to form biofilms compared to the parental strain (****P<0.0001, one-way ANOVA with Tukey’s *post hoc* correction) and a 40% attenuation when compared to the *A909:rev* control (*P<0.05, one-way ANOVA with Tukey’s *post hoc* correction).

We also used scanning electron microscopy to image biofilms on glass coverslips.

The cell morphology of the Δ*sip* strains looked similar, overall, to the *rev* and WT comparators, forming chains of several dozen dividing cocci. However, while the *rev* strains adhered to the cover slip surface and aggregated in robust biofilms, the Δ*sip* strains did not organize into biofilm aggregates (**Fig. 6**).

**Figure 6:**
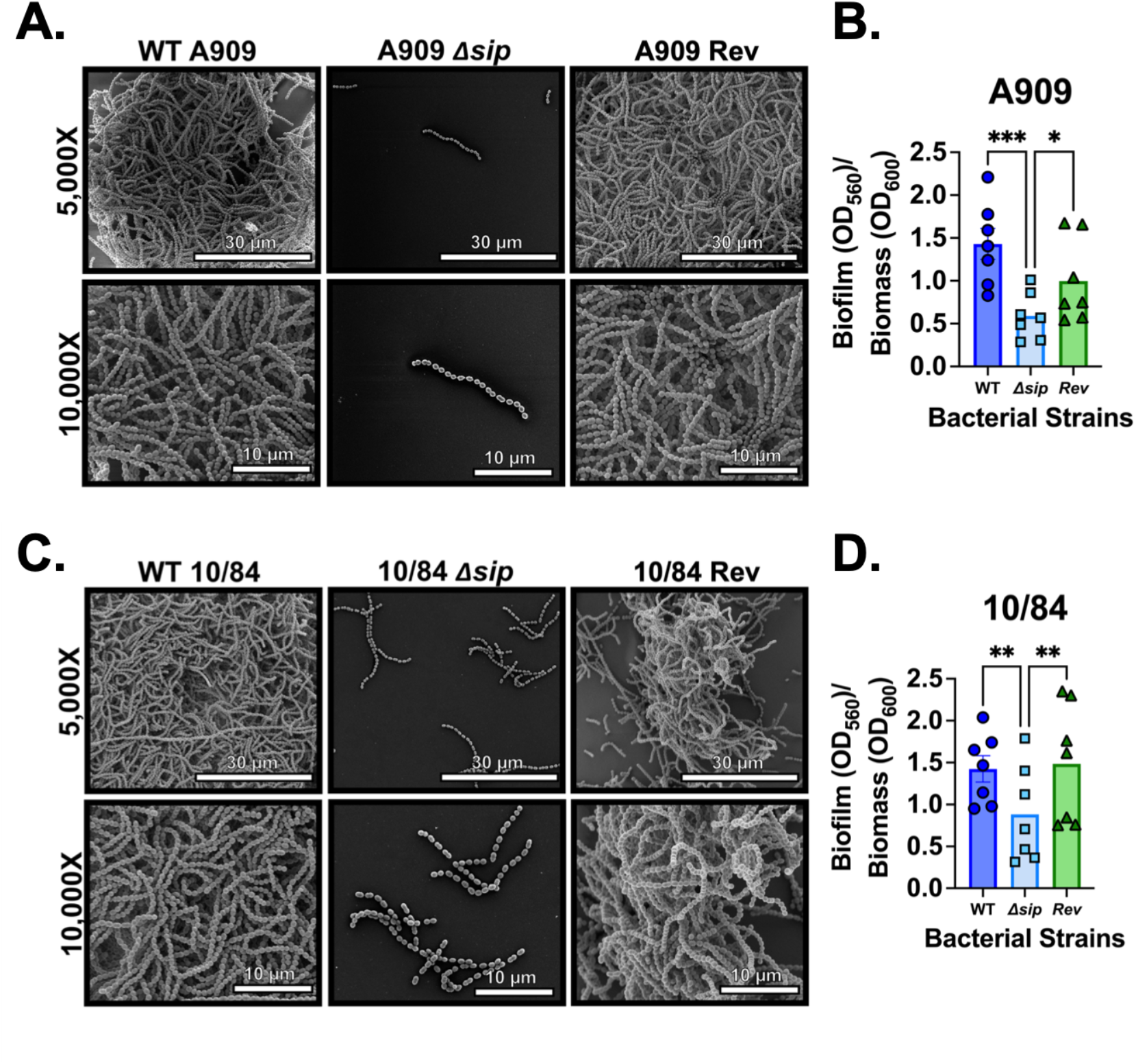
Sip contributes to biofilm formation by GBS. High resolution field-emission gun scanning electron microscopy analysis of wild-type (WT) A909 (A) reveals large tertiary architectural structures of cells indicative of robust biofilm formation. Conversely, the isogenic *Δsip* mutant adheres sparsely to the abiotic surface without forming tertiary structured biofilms, a result that was reversed in the *Rev* control strain. Quantification of biofilms by spectrophotometric analysis indicates WT and *Rev* A909 forms significantly more quantifiable biofilm than the isogenic *Δsip* (B, each datapoint represents an independent biological replicate; n=8 independent biological replicates, each with 3 technical replicates). The same biofilm electron micrograph (C) and biofilm quantification (D; n=8 independent biological replicates, each with 3 technical replicates ) patterns were observed in the CNCTC 10/84 background strain. \**p<0.05*, \*\**p<0.01, ***p<0.001*, one-way ANOVA with Tukey’s *post hoc* correction.

### GBS Δsip mutants show decreased attachment to and penetration of ex vivo human fetal membranes

To examine whether the altered surface characteristics that decreased biofilm formation by Δ*sip* GBS would change these strains’ association with human tissue relevant to perinatal infection, we imaged capsule-stained adhered and penetrating GBS in cross sections of freshly collected human fetal membranes. We also performed adhesion assays based on CFU counts recovered following experimental fetal membrane exposure to our GBS variants.

We found decreased association between GBS Δ*sip* strains and fetal membranes by both measures, compared to *rev* and WT controls. There was reduced GBS staining intensity on cross sections of fetal membranes infected with Δ*sip* strains, compared to WT and *rev* controls. In adhesion assays, CFU counts from Δ*sip* infected membranes were 1-2 log-fold less than WT and *rev* controls, differences that were significant by one-way ANOVA testing with Tukey’s *post hoc* correction (**Fig. 7**).

**Figure 7:**
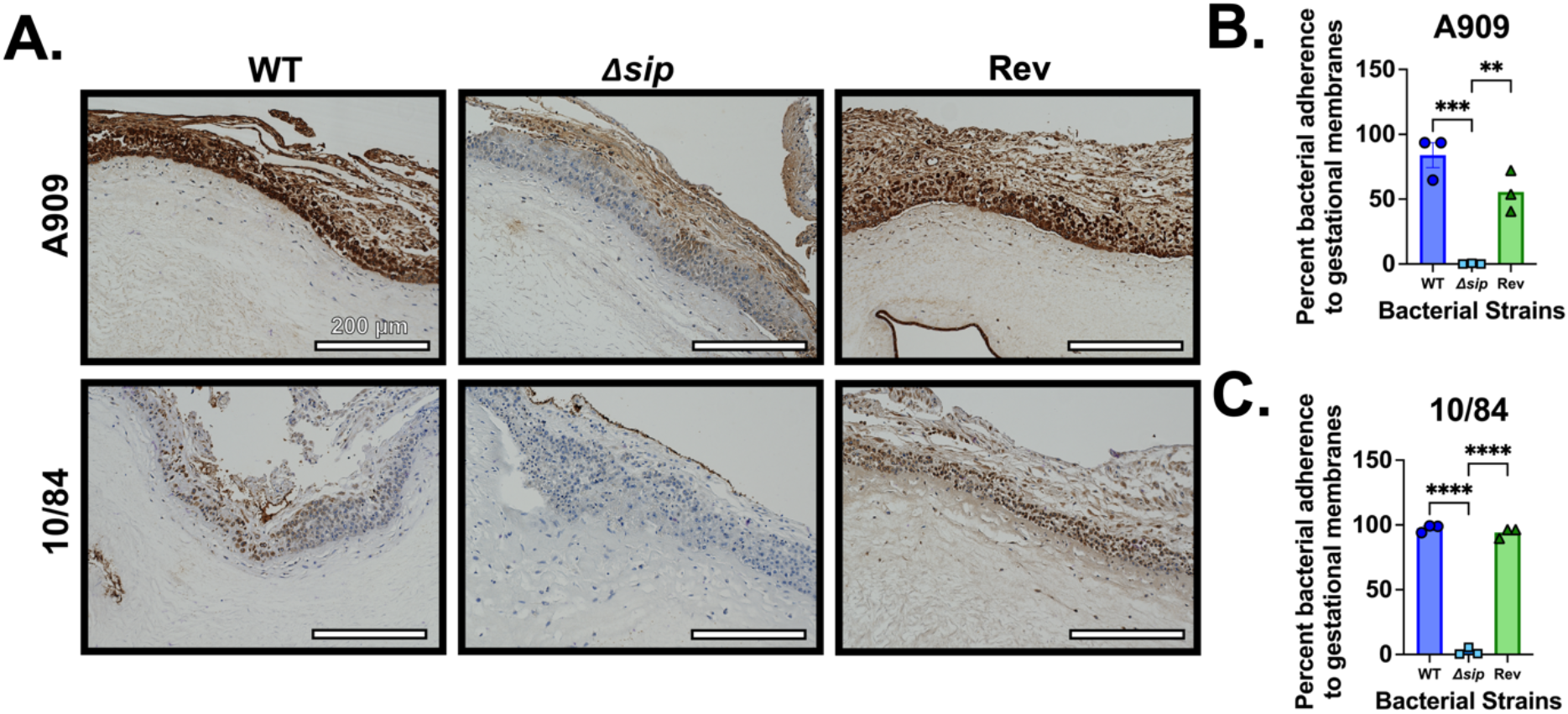
Sip contributes to adherence to primary human fetal membrane explants. Primary human fetal membrane samples were colonized on the choriodecidual surface and co-incubated with GBS WT, Δ*sip*, and *Rev* strains prior to being fixed, sectioned, and stained with a rabbit polyclonal anti-GBS antibody. Significant differences were observed microscopically, with decreased association between the Δ*sip* strains and the fetal membranes compared to WT and *Rev* controls (**A**, representative CNCTC 10/84 and A909 background images). Quantitative culture results of homogenized membrane samples prior to fixation showed a significant decrease in Δ*sip* adherence in both A909 (**B**) and 10/84 (**C**) backgrounds. Statistical testing by ANOVA with Tukey’s *post hoc* correction where each datapoint represents one independent biological replicate; \*\**p<0.01*, \*\*\**p<0.001*, \*\*\*\**p<0.*0001.

### GBS Δsip mutants have impaired uterine invasion in a mouse model of vaginal colonization and ascension

Given the altered interactions we observed between Δ*sip* GBS and cell models of innate immunity, and the adhesion, biofilm, and tissue persistence defects presented by the mutants, we sought to understand if these mutant phenotypes led to altered reproductive tract colonization characteristics *in vivo*. We used a nonpregnant female C57BL/6J mouse model in which eight-week old estrus-synchronized mice underwent standardized vaginal colonization with overnight cultures of CNCTC 10/84:Δ*sip* or CNCTC 10/84:*rev*. Following colonization and a 48-hour equilibration period, daily vaginal swabs were used to make PBS suspensions, which were then plated for CFU quantification on GBS-specific chromogenic agar. At the end of seven days of daily swabs, the colonized mice were sterilely dissected for isolation of cervical and uterine tissue, which was homogenized and plated for GBS CFU quantification.

We observed no significant differences in vaginal colonization density over the seven-day swab phase (one-way ANOVA with Tukey’s correction; each day’s swab CFU density was compared between WT and Δ*sip*), nor were there significant differences in rates of colonization clearance between the two GBS variants (**Fig. 8A**). We did observe differences in uterine CFU density at the end of the week, however, with increased uterine GBS burden in the WT strain compared to Δ*sip* (*p<0.05, Mann-Whitney U test, **Fig. 8B**). No difference was measured in cervical CFU density between the two strains. The uterine result is consistent with a GBS-host interaction model in which Sip has an innate immune suppressive effect, promoting persistence of WT GBS in the uterus after vaginal colonization, while Δ*sip* leads to increased inflammasome activation, increased cytokine signaling, and more efficient clearance from the immunologically protected uterine compartment.

**Figure 8:**
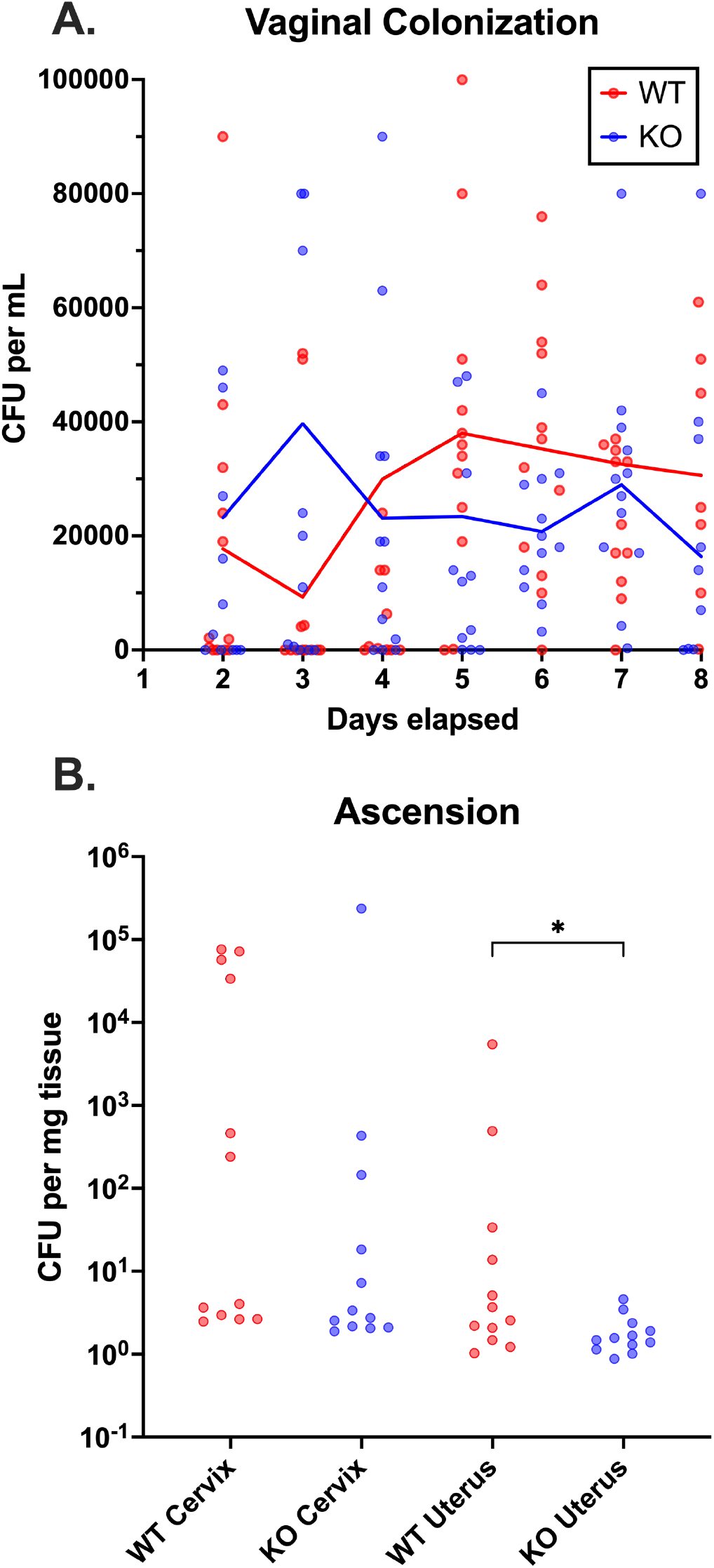
Sip affects uterine ascension but not vaginal colonization or cervical ascension. WT C57BL/6J mice (n=12) were vaginally colonized and swabbed daily from day two through eight post-colonization (**A**, solid lines indicate mean values for each day; no statistical difference by Mantel-Cox testing). On day 8, the animals were sterilely dissected for cervix and uterine tissue, which was used for quantitative GBS culture (**B**). Each datapoint represents one mouse; data lines indicate median (\**p<0.05*, Mann-Whitney).

## Discussion

To our knowledge, this is the first report of using CRISPRi in GBS to study a large gene set for roles in pathogenesis. The main advantage of CRISPRi is that generation of specific gene knockdowns is faster than targeting loci for chromosomal deletion, and—in our experience—less prone to unintended outcomes. Problems such as off-target crossover events by a mutagenesis plasmid or unwanted reversion to the WT genotype at the final plasmid excision step can substantially hinder efforts at chromosomal recombination but are not factors when using CRISPRi.

Another appealing aspect of CRISPRi is that it can be used as an approach to study the functions of essential or conditionally essential genes. Partial suppression of essential gene transcription can result in growth and morphotype alterations, which may render essential gene knockdown strains not directly comparable to WT or sham-targeted WT equivalent variants. However, this is a preferable situation to having only chromosomal deletion approaches available, which offer few options for studying essential gene function. Two genes in the CRISPRi knockdown collection for this investigation were predicted to be essential based on previous Tn-seq analysis (W903_RS00250 and W903_RS05170)^64^; both were transcriptionally suppressed based on RT-qPCR data and included in our analysis pipeline.

We used CRISPRi to query a curated set of genes that encode predicted surface- trafficked proteins, based on the presence of N-terminal signal peptide sequences. We were not fully successful in generating CRISPRi knockdown variants targeting all 75 signal peptide-encoding genes in the CNCTC 10/84 genome, nor were we able to make two knockdown strains for all 70 of the genes we targeted, as 19 genes were only targeted with a single sgRNA. This points to some drawbacks of CRISPRi. Cloning can still fail in the *E. coli* or GBS transformation stages of CRISPRi, for reasons that are not necessarily clear.

Our extensive RT-qPCR dataset for our collection of GBS knockdown strains also highlights the variability of gene expression knockdown by CRISPRi. We measured expression changes in our target genes that ranged from no suppression to 2^-12^ relative to a non-targeted dCas9 comparator. Depending on a gene’s function and baseline expression profile, partial knockdown could lead to a significantly altered phenotype, a modestly altered phenotype, or no significant change. This fact points to the importance of using multiple sgRNA protospacers to target each gene of interest and highlights that CRISPRi is most appropriately used as a screening approach to identify genetic targets for more definitive study by formal chromosomal deletion.

When we performed macrophage-like THP-1 cell coincubation with GBS variants from our CRISPRi collection, we were surprised to find a range in TNF-α and IL-1β secretion responses that spanned from suppressed to significantly exaggerated. Our hypothesis had been that removing GBS surface proteins would tend to decrease antigenic signaling to phagocytes, resulting in less pro-inflammatory cytokine signaling. The fact that a substantial number of our CRISPRi variants led to increased pro-inflammatory cytokine release suggests that innate immune repression may be an important summative influence of multiple GBS surface proteins. Innate immune suppression is a known function of the GBS sialylated capsule^16,65,66^, and multiple previously studied GBS surface-trafficked proteins are known to promote GBS infection by evading immune activation. For example, GBS pili have been shown to partially block phagocytic killing by macrophages and neutrophils through resistance to antimicrobial peptide-mediated killing^67^; the C5a peptidase protein encoded by the *scpB* gene decreases neutrophil attachment to GBS by inactivating complement factors that threaten survival during bloodstream invasion^24,25^; and the SHP/RovS system is an intercellular communication system shown to enable GBS populations to respond in a coordinated manner to molecular threats such as might arise from innate immune activation during infection^68^.

Among the surface-trafficked proteins we examined, knockdown of the gene that encodes Sip had the greatest effect on IL-1β release from co-incubated THP-1 cells, with *sip* CRISPRi knockdown leading to a 3.5-fold increase in IL-1β secretion. Sip targeting also had a significant effect on THP-1 secretion of TNFα, increasing it 2.4-fold over the nontargeted control strain coincubation. Sip has been the focus of attention as a novel GBS vaccine candidate or as a GBS vaccine adjuvant for decades. Sip recently received new attention in an analysis of surface proteins detected among a large South African cohort of GBS isolates, among which it was noted to be a top vaccine target candidate based on its conserved expression and high antigenic potential according to bioinformatic modeling^69^ . First described in 2000 as a highly conserved GBS surface protein with cross-serotype protective properties when purified recombinant protein preparations (rSip) were administered prior to an animal model of GBS sepsis^38^, little has been published on Sip biology *in situ* on the GBS external surface. In studies of rSip as a possible protein adjuvant for a GBS (or other bacterial) vaccine, the purified protein has been shown to stimulate toll-like receptors (TLR) 2 and 4^40,70^. While this finding does not align neatly with our finding of increased pro-inflammatory cytokine expression when THP-1 cells were exposed to GBS lacking Sip, differences in host cell response may be influenced by the contextual presentation of GBS surface protein antigens. In other words, Sip displayed on the outer surface of GBS cells may dampen pro-inflammatory pathway activation whereas purified protein may have an immune stimulatory effect.

Our subsequent investigations of Sip function using targeted deletion strains in CNCTC 10/84 and A909 backgrounds demonstrated additional roles that Sip may play in GBS disease pathogenesis. Upon isolation of the Δ*sip* mutants, we noticed evidence of altered surface characteristics, including apparent decreased aggregation in biofilms when grown *in vitro*, an observation that we quantified and expanded upon in experiments demonstrating decreased association with human fetal membrane explants. Past work has demonstrated that Sip is accessible to antibodies in representative strains from all ten known capsular serotypes and—based on transmission electron microscopy of gold-conjugated secondary antibodies binding anti-Sip antibody—localizes to the cleavage planes and distal poles of GBS cells^39^. We interpret our data, in the context of previous studies, as indicating that decreased expression of Sip at these sites either reduces interbacterial and bacterial-host adhesion directly or affects the function of different proteins that promote surface interactions.

The imputed protein structure of Sip does not indicate an obvious biological function, much less important roles in immune evasion and adhesion. Conserved domains include a lysin motif (LysM), whose 44 amino acids are predicted to promote protein-peptidoglycan binding. LysM-peptidoglycan interactions can underlie various mechanistic functions, but a common final pathway of LysM activity is pattern-specific peptidoglycan hydrolysis^71^. Localization of GBS Sip to the cleavage plane suggests a possible role for cell wall hydrolysis promoting GBS division. A significant portion of the protein is identified as a possible ribonuclease E motif, based on amino acid sequence signatures, although the predicted folding of this portion of the protein is low-confidence and its role at the GBS cell surface unclear.

An examination of partial protein homology in other *Streptococcal* species uncovered a report describing a LysM-containing surface protein in *Streptococcus suis* with 41% identity to the GBS protein. Like the GBS protein, a recombinant preparation of the *S. suis* factor conferred partial resistance to experimental *S. suis* infection in a mouse model. Furthermore, a Δ*lysM S. suis* strain showed reduced virulence compared to wild type and a plasmid-complemented strain, and the mutant was more susceptible in a whole-blood killing assay, suggesting that the intact protein may share the immune evasion roles suggested by our experiments using GBS^72^.

The near-complete conservation of the *sip* gene across GBS strains, its demonstrated potential as a vaccine component, and the immunomodulatory properties of the surface protein *in situ* demonstrated here make Sip an important GBS molecule for additional experimental study. Open questions about its role on the GBS cell surface, how it contributes to suppression of cytokine secretion by macrophage-like cells, and what functions it may serve during invasive disease—other than potentially promoting persistence in the pregnant uterus as indicated by mouse experiments in this study—are key topics for consideration and will be the focus of future effort by our group.

## Methods

### Ethics statement

Animal experiments were performed under an approved IACUC protocol (#23012501) at University of Pittsburgh. Collection of human fetal membranes from non-laboring cesarian section deliveries was conducted under Institutional Review Board approval with informed consent from the Vanderbilt University IRB (#181998).

### Reagents

RPMI 1640 +L-Glutamine, BD Bacto dehydrated tryptic soy broth (TSB), Luria-Bertani (LB) medium, erythromycin, DNA-free DNA removal kit, TRIzol reagent, penicillin-streptomycin-glutamine (PSG) 100×, antibiotic (penicillin-streptomycin)-antimycotic (amphotericin) solution, PBS, DPBS +CaCl2 +MgCl2 (DPBS^+/+^), TNF-α FAM-MGB primer/probe (Hs01113624_g1), IL-1β FAM-MGB primer/probe (HS01555410_m1), Bio-Rad iTaq Universal Sybr Green One-Step, MagMAX Viral/Pathogen Ultra Nucleic Acid kit, and GAPDH FAM-MGB primer/probe (Hs02758991_g1) were purchased from Thermo-Fisher (Waltham, MA). THP1-Blue cells were purchased from InvivoGen (San Diego, CA).

Charcoal-stripped and dextran-treated fetal bovine serum (FBS), TNF-α ELISA kit and IL-1β ELISA kits were purchased from RCD Systems (Minneapolis, MN). Ethyl alcohol, non-enzymatic cell dissociation solution, and phorbol 12-myristate 13-acetate (PMA) were purchased from Sigma-Aldrich (St. Louis, MO). RNeasy mini kit was purchased from Qiagen (Germantown, MD). SsoAdvanced universal supermix and iScript cDNA synthesis kit were purchased from BioRad Laboratories (Hercules, CA). The irreversible caspase-1 inhibitor, Z-YVAD-FMK, was purchased from Abcam (Waltham, MA). QIAprep spin miniprep kits were purchased from Qiagen (Hilden, Germany). Rabbit polyclonal anti-GBS antibody was purchased from Abcam (Cambridge, UK).

### Bacterial strains and growth conditions

GBS strains A909 (serotype Ia, sequence type 7) and CNCTC 10/84 (serotype V, sequence type 26) and their derivatives were grown at 37°C (or 28°C when the temperature-sensitive pMBsacB plasmid was present and extrachromosomal) under stationary conditions in TSB supplemented with 5 µg/ml erythromycin as needed for selection. *E. coli* DH5α for cloning were purchased in chemically competent preparations from New England Biolabs, transformed according to manufacturer instructions, then grown at 37°C (or 28°C with extrachromosomal pMBsacB present) with shaking in LB medium supplemented with 300 µg/ml erythromycin for plasmid propagation.

### Identification of conserved signal peptide-encoding genes

Conserved GBS signal peptide-encoding genes in CNCTC 10/84 were identified using the publicly accessible bacterial genome dataset maintained by the United States Department of Energy Joint Genome Institute’s (JGI) Integrated Microbial Genomes and Microbiomes System (IMG/M; https://img.jgi.doe.gov/m/). First, we performed a gene search querying the CNCTC 10/84 genome (IMG/M taxon ID 2627854227) and using the [is signal peptide = yes] designator. We saved the resulting set, then searched the genome database for *Streptococcus agalactiae* genomes, saving this set as a searchable collection.

To cross-reference the set of CNCTC 10/84 signal peptide-encoding genes against the set of GBS genomes, we used the Profile & Alignment tool in the IMG/M “Gene Cart” menu. The maximum E-value was set to 0.1 and the minimum percent identity was set to 10 percent. The process was repeated, increasing the percent identity by 10-percent increments to 90 percent. With each iteration, we saved the output table indicating which genes in the set exceeded the identity threshold. Once we had generated tables for each 10-percent threshold, we tallied—for each gene—the maximum percent identity recorded. This gave us a quantifiable measure of conservation for each gene in the set.

### Creation of the surface-trafficked protein CRISPRi library

For each of the 66 conserved surface-trafficked CNCTC 10/84 genes, two targeting CRISPRi protospacers were designed using the Broad Institute’s CRISPick server, using *S. pyogenes* PAM settings given the homology between groups A and B *Streptococcus* CRISPR/Cas9 mechanisms. Full-length gene sequences were entered onto the server for each gene, generating lists of potentially active targeting sites in the coding sequences.

Protospacers were selected based on complementarity to the antisense strand of the target gene and location, whenever possible, in the first third of the coding sequence.

Once the protospacer set was determined, custom forward and reverse protospacer oligonucleotide preparations were obtained from Integrated DNA Technologies (IDT). The oligonucleotides were designed so that, once annealed, the resulting double-stranded construct would have *BsaI* restriction site-compatible sticky ends to permit cloning into the sgRNA expression plasmid p3015b as previously described^35^. Cloning and transformation of chemically competent DH5α *E. coli* was followed by selection for erythromycin resistance and visible expression of red-tinted mCherry fluorescent protein, encoded as a marker on p3015b. Putative successful transformants were screened with colony PCR using the forward protospacer oligonucleotide as one primer and a conserved reverse oligonucleotide complementary to the p3015b plasmid, upstream of the protospacer insertion site, as the second primer.

Successful cloning was determined based on the presence of a 1000-bp band on a standard agarose electrophoresis gel.

Plasmid minipreps were performed on overnight cultures of successful *E. coli* clones using the Qiagen QIAprep Spin Miniprep Kit according to manufacturer instructions. Purified p3015b with targeting protospacer was then used to transform electrocompetent CNCTC 10/84:*dCas9* using established techniques^32,73,74^. Putative GBS knockdown strains were selected on TSB with erythromycin with confirmatory observation of mCherry expression, then stored as frozen glycerol stocks.

### RT-qPCR of CRISPRi target gene expression

Primers for RT-qPCR screening of CRISPRi target gene expression were designed and ordered on the IDT website using the PrimerQuest tool. We used primers optimized for Bio-Rad iTaq Universal Sybr Green One-Step reagents. RNA was extracted using MagMAX Viral/Pathogen Ultra Nucleic Acid kit reagents, according to manufacturer instructions, from cultures of the GBS CRISPRi library strains grown overnight in sterile deep-well 96- sample plates. The extraction was performed with a Hamilton Nimbus robotic liquid handling instrument with an inset Thermo Presto magnetic bead purification device.

Each CRISPRi strain RNA sample was extracted in reverse transcriptase-containing and reverse transcriptase-negative master mixes. Following extraction, the RNA samples were DNA depleted using Thermo DNase and inactivation agent (Cat. # AM1906) according to manufacturer instructions except that the 37°C incubation was allowed to proceed for 90 minutes (rather than 30 minutes). DNA depletion was tested by comparing reverse transcriptase-positive and -negative RT-qPCR curves.

RT-qPCR testing was performed using a Bio-Rad CFX96 Touch real-time PCR thermocycler set to 40 cycles with temperature settings in accordance with the iTaq Universal Sybr Green One-Step reagent instructions. Gene expression quantification was calculated using the Livak method^75^ with normalization to the GBS *recA* gene^76^ as an endogenous control and a CNCTC 10/84:*dCas9* strain with the p3015b plasmid lacking a targeting protospacer as a WT-equivalent baseline expression comparator.

### Targeted deletion of Sip genes

The *sip* genes in CNCTC 10/84 and A909 were deleted with the temperature- and sucrose-sensitive plasmid pMBsacB, using previously described techniques^32^.

Approximately 500-bp upstream and downstream homology arms were amplified from the respective chromosomes and cloned into the modular restriction enzyme sites on pMBsacB such that chromosomal insertion and subsequent excision of the plasmid would result in either a markerless deletion of *sip* or reversion to the WT genotype. After the final sucrose counterselection step against retention of the plasmid sequence, PCR across the *sip* site on the chromosome was used to identify putative Δ*sip* and WT reversion strains.

Whole genome sequencing of chromosomal DNA from the different strains was performed prior to their use in disease modeling experiments.

### Ethanol killing of GBS

GBS strains were grown in 50 mL of TSB at 37°C without shaking. GBS were then ethanol-killed using a slightly modified protocol from that described in work published by Lu et al^77^. Briefly, cultures of GBS were washed twice by centrifugation with cold DPBS^+/+^ and resuspended in 5 mL of cold DPBS^+/+^. Cultures were serially diluted onto blood agar to determine concentration (CFU/mL). 100% ethanol was added in equal increments over 15 minutes to a final concentration of 70% at 4°C with gentle rocking. GBS was rocked at 4°C for an additional hour. GBS was washed twice with DPBS^+/+^ as before and resuspended in DPBS^+/+^. Cells were tested for viability by spotting onto blood agar and inoculating into THB. Ethanol-killed GBS (GBS^EK^) were aliquoted and stored at -80°C avoiding freeze/thaw cycles.

### Growth and PMA treatment of THP1-Blue cells

THP-1 Blue cells were grown and passaged in RPMI containing 10% FBS, 1% PSG and 100 µg/mL normocin (referred to as RPMI^+/+/+^ media) and treated with PMA to a final concentration of 5 ng/mL overnight at 37°C with 5% CO2 to mature into macrophage-like cells. Cells were collected using non-enzymatic dissociation solution at 37°C with 5% CO2 for 5 min followed by gentle scraping.

### THP1-Blue stimulation for cytokine analysis

PMA-treated THP1-Blue cells were plated at 400,000 cells/well in RPMI (lacking both FBS and antibiotic; RPMI^-/-^) in triplicate for each condition into a 96-well tissue culture dish and rested for 30-90 minutes at 37°C with 5% CO2. Media was aspirated and 150 µL of fresh RPMI^-/-^ was added to wells before GBS^EK^ strains were added at a multiple of infection (MOI) of 50:1. In some instances, PMA-treated THP-1 Blue cells were pretreated with 10 µM of the irreversible caspase-1 inhibitor Z-YVAD-FMK for 1 hr prior to stimulation with GBS . All stimulated macrophages were incubated at 37°C with 5% CO2 for 22-24 hr. Following stimulations, culture media in wells (technical replicates pooled) were centrifuged at 4°C and 13,000 RPM for 3 min in a tabletop micro-centrifuge. Supernatants were stored at - 80°C until cytokine analysis by ELISA.

### THP-1 Blue stimulation for qRT-PCR analysis

PMA-treated THP-1 Blue cells were plated at 5×10^6^ cells/well in RPMI^-/-^ into a 6-well tissue culture plate and rested for 30 min. Media was aspirated and 1 mL of fresh RPMI^-/-^ was added to wells before GBS^EK^ strains were added at MOI 50:1 for 4 hr at 37°C with 5% CO2. Total RNA was isolated using TRIzol and scraping each well with a flat blade cell lifter and stored at -80°C. RNA was extracted from TRIzol suspension per the manufacturer’s instructions. RNA quantity and quality was determined using a NanoDrop before being treated with DNase as described by the manufacturer. Next, 1 µg of cDNA was synthesized using the Applied Biosystems ProFlex PCR system. Finally, 2 µL of cDNA was subject to real-time q-PCR using SsoAdvanced Universal Supermix with a 20 µL total reaction volume using an Applied Biosystems QuantStudio 3 thermocycler. All samples were run in triplicate and data was analyzed using the ΔΔCt method.

### Cytokine analysis by ELISA

Cytokines from macrophage stimulation experiments were analyzed by ELISA per the manufacturer’s kit instructions for human TNF-α and IL-1β.

### Quantitative analysis of biofilms

Biofilm formation was quantified by crystal violet staining of overnight static cultures as previously described^15^. Briefly, GBS cultures were grown overnight in Todd-Hewitt broth (THB) and sub-cultured at 1:100 into 100 µL fresh THB +1% glucose in 96-well culture plates. Cultures were incubated statically at 37°C in ambient air overnight. The following day, OD600 was measured evaluate cell density, and cultures were decanted and washed three times before staining with 1% crystal violet. Wells were washed three times with water and allowed to dry before crystal violet was re-solubilized in 80% ethanol: 20% acetone solution and the total biofilm quantification was measured at OD560. Total biofilm to biomass was calculated as the ratio of OD560 of re-solubilized crystal violet to the OD600 measurement of total cell density.

### Bacterial co-culture assays on explant human fetal membranes

Human placenta and fetal membranes were isolated from term, healthy, non-laboring caesarean section procedures. Fetal membranes were separated from the organ and 12 mm diameter tissue pieces were cut with a sterile biopsy punch. Tissue pieces were cultured amnion side down in modified Roswell Park Memorial Institute medium 1640 supplemented with L-glutamine, HEPES, 1% fetal bovine serum (mRPMI 1640), and antibiotic/antimycotic mixture (Gibco, Carlsbad, California). Tissues were incubated overnight at 37°C in room air containing 5% CO2. The following day, tissues were washed 3 times sterile phosphate buffered saline (pH 7.4) and placed again in mRPMI 1640 lacking the antibiotic/antimycotic supplement. Bacterial cells were added at a final concentration of 1 × 10^7^ cells per mL to the choriodecidual surface of the fetal membranes. Uninfected membrane tissues were also maintained as a negative control. Co-cultures were incubated at 37°C containing 5% CO2 for 24 hours. After co-incubation, a portion of each membrane sample was separated, weighed, homogenized, and plated on solid agar media for CFU quantitation; the remainder or each sample was used for fixation and immunohistochemical analysis.

### Immunohistochemical analysis of bacterial association with human fetal membranes

Samples were fixed in neutral buffered 10% formalin before being imbedded in paraffin blocks. Tissues were cut into 5µm thick sections and multiple sections were placed on each slide. Samples were washed with xylene for 2 minutes. Heat-induced antigen retrieval was performed using Epitope Retrieval 2 solution (Leica Biosystems) for 20 min. Slides were stained with a 1:100 dilution of the rabbit polyclonal anti-GBS antibody (ab78846; Abcam) for 1 hour. The Bond Polymer Refine detection system (Leica Biosystems) secondary detection system was applied. Slides were counter-stained with eosin, alcohol dehydrated, and mounted with glass coverslips before light microscopy was performed with an EVOS light microscope.

### Field emission gun scanning electron microscopy (FEG-SEM) analysis

Samples were prepared for scanning electron microscopy analyses as previously described. Briefly, samples were fixed in 2.0% paraformaldehyde, 2.5% glutaraldehyde, in 0.05 M sodium cacodylic acid overnight at room temperature. The following day, samples were sequentially dehydrated with increasing concentrations of ethanol (25%, 50%, 75%, 95%, and 100%) for 1 hour each step. Samples were dried at the critical point using a carbon dioxide critical point dryer (Tousimis) prior to mounting on aluminum SEM stubs and plasma sputter coating with approximately 20 nm of 80/20 gold/palladium. Sample edges were painted with colloidal silver to facilitate charge dissipation and imaged with an FEI Quanta 250 field-emission gun scanning electron microscope. ^1^. Briefly, samples were fixed with 2.5% glutaraldehyde, 2.0% paraformaldehyde, in 0.05 M sodium cacodylate buffer (pH 7.4) at room temperature for 24 hours. Subsequently, samples were washed three times with 0.05 M sodium cacodylate buffer and sequentially dehydrated with increasing concentrations of ethanol. After dehydration, samples were dried with a Tousimis CO2 critical point dryer, mounted onto aluminum stubs, and painted at the sample edge with colloidal silver to dissipate excess charging. Samples were imaged with an FEI Quanta 250 field emission gun scanning electron microscope at an accelerating voltage of 5.0 KeV at 5,000X to 10,000X magnification.

### Mouse model of vaginal colonization and ascending chorioamnionitis

Single-housed 8-week old, female C57BL/6J mice were used in an established vaginal colonization and ascending infection model^9^ with minor protocol changes.

Following two days of estrus synchronization with 0.5 mg subcutaneous β-estradiol, CNCTC 10/84 Δ*sip* or *rev* strains was grown overnight in 5 mL TSB, pelleted by centrifugation the next morning, then resuspended in a 1:1 sterile PBS and 10% gelatin mixture. Mice were vaginally colonized with 50 μL of this mixture, then returned to their cages. After a 48 hr equilibration period, the mice were vaginally swabbed with a moistened sterile nasopharyngeal swab, which was then swirled three times in 300 μL sterile PBS. This swab resuspension was then serially diluted and plated on GBS-specific chromogenic agar plates for next-day CFU enumeration.

At the end of seven days of swabbing, the mice were euthanized and dissected for sterile removal of the cervix and uterus. These tissue samples were weighed, homogenized, and plated on chromogenic agar for CFU enumeration.

## Figures & statistical analysis

Except where otherwise noted, experiments were performed on independent biological replicates with triplicate technical replicates. Technical replicate values were averaged, and statistical analyses were performed on biological replicate means. Figures were generated in Graphpad Prism and heatmapper.ca. Statistical analyses were performed in Graphpad Prism and Python (v 3.10.4) with the SciPy library (v 1.8.1).

## Disclosures

The authors have no financial or legal conflicts of interest to report.

## Acknowledgments

This work was supported by the National Institutes of Health (NIH) R21AI178067 (T.A.H. and D.M.A.), R01AI182835 and R01AI177991 (T.A.H.), R01HD090061 and R01HD113675 (J.A.G.), the March of Dimes #6-FY24-0009 (J.A.G.), and the Burroughs Wellcome Fund Next Gen Pregnancy Initiative # 1275387 (J.A.G.), and Department of Veterans Affairs Merit Award I01BX005352 Office of Research (J.A.G). This work was also supported by NIH R01AI134036 (J.A.G. and D.M.A.) and by the Rise-Up Summer Research Program (Office of Research and Development, Department of Veterans Affairs, I01BX007005). The content of this manuscript is solely the responsibility of the authors and does not necessarily represent the official views of our funders.

## Supplemental data

**Supplemental Data 1: Tab 1 (CNCTC Signal Peptide Genes)** Data table listing the set of CNCTC 10/84 signal peptide-containing genes that were also present among a subset of the 654-genome screening collection. The column labeled “Conserved” indicates those 66 genes that were in the CRISPRi library. **Tab 2 (Conserved Knockdown Library)** Lists the knockdown strains in the CRISPRi library. The protospacer number in the “Gene- Protospacer ID” column indicates where in the gene coding sequence dCas9 was targeted for that knockdown strain. **Tab 3** (**RT-qPCR)** shows normalized expression data from the CRISPRi library strains. **Tab 4 (CRISPRi Cytokine Profiling)** shows ELISA results from THP-1 macrophage coincubation with ethanol-killed strains from the CRISPRi library.

## Notes

### Competing Interest Statement

The authors have declared no competing interest.

